# Chronic innate immune impairment and ZIKV persistence in the gastrointestinal tract during SIV infection in pigtail macaques

**DOI:** 10.1101/2024.08.23.609309

**Authors:** Jennifer Tisoncik-Go, Thomas B. Lewis, Leanne S. Whitmore, Kathleen Voss, Skyler Niemeyer, Jin Dai, Paul Kim, Kai Hubbell, Naoto Iwayama, Chul Ahrens, Solomon Wangari, Robert Murnane, Paul T. Edlefsen, Kathryn A. Guerriero, Michael Gale, Deborah H. Fuller, Megan A. O’Connor

## Abstract

Mosquito borne flaviviruses, including dengue (DENV) and Zika (ZIKV) viruses, have caused global epidemics in areas with high HIV prevalence due to the expanded geographic range of arthropod vectors. Despite the occurrence of large flavivirus outbreaks in countries with high HIV prevalence, there is little knowledge regarding the effects of flavivirus infection in people living with HIV (PLWH). Here, we use a pigtail macaque model of HIV/AIDS to investigate the impact of simian immunodeficiency virus (SIV)-induced immunosuppression on ZIKV replication and pathogenesis. Early acute SIV infection induced expansion of peripheral ZIKV cellular targets and increased innate immune activation and peripheral blood mononuclear cells (PBMC) from SIV infected macaques were less permissive to ZIKV infection *in vitro*. In SIV-ZIKV co-infected animals, we found increased persistence of ZIKV in the periphery and tissues corresponding to alterations in innate cellular (monocytes, neutrophils) recruitment to the blood and tissues, decreased anti-ZIKV immunity, and chronic peripheral inflammatory and innate immune gene expression. Collectively, these findings suggest that untreated SIV infection may impair cellular innate responses and create an environment of chronic immune activation that promotes prolonged ZIKV viremia and persistence in the gastrointestinal tract. These results suggest that PLWH or other immunocompromised individuals could be at a higher risk for chronic ZIKV replication, which in turn could increase the timeframe of ZIKV transmission. Thus, PLWH are important populations to target during the deployment of vaccine and treatment strategies against ZIKV.

**Author Summary:** Flaviviruses, including Zika virus (ZIKV), cause global epidemics in areas with high HIV prevalence. Yet questions remain as to whether ZIKV disease is altered during an immunocompromised state and the potential immune mechanisms contributing to enhanced disease. This is essential to our understanding of ZIKV disease in people living with HIV (PLWH). Here, we use a non-human primate (NHP) model of HIV/AIDS to investigate the impact of immune suppression on ZIKV replication and pathogenesis. The use of the NHP model was critical for the assessment of longitudinal specimens across tissues that are active sites of flavivirus replication and host immune responses. This study broadly demonstrates that ZIKV pathogenesis is altered and more persistent in states of immunosuppression. Collectively, this study suggests that in PLWH and immunocompromised individuals, other arboviruses, including dengue and West Nile viruses, could similarly alter pathogenesis and/or viral peristance in tissues. Furthermore, this study highlights the need to prioritize immunocompromised individuals in the design and rollout of vaccines against arboviral diseases.

## Introduction

Flaviviruses, including dengue (DENV), West Nile (WNV), yellow fever (YFV), and Zika (ZIKV) viruses, are single-stranded RNA viruses that are transmitted to people through the bite of an infected mosquito and have caused global epidemics in recent years. Flavivirus infection commonly causes mild clinical manifestations; however, more severe hemorrhagic or encephalitic disease can occur and the mechanisms underlying severe disease are not fully understood. Furthermore, vulnerable populations including children, pregnant women, and immunocompromised individuals, including people living with HIV (PLWH) may be at a higher risk for more severe flavivirus disease. Currently, highly effective vaccines against DENV, WNV, and ZIKV do not exist. While the live-attenuated YFV vaccine 17D is available, it is contraindicated in infants and in immunosuppressed individuals and is relatively contraindicated in the elderly, pregnant women, and PLWH due to poor immunogenicity or severe adverse reactions(1–4). Therefore, there is a need to better understand flavivirus pathogenesis in at risk populations.

ZIKV transmission is primarily via mosquito bite; however, transmission also occurs through sexual intercourse and from mother to fetus(5–7). ZIKV infection usually results in mild and self-limiting symptoms, but it is also associated with the neurological disorder Guillain-Barré syndrome (GBS) in adults and congenital Zika syndrome (CZS) during *in utero* exposure(8, 9). CZS is characterized by severe defects in cranial morphology, ocular abnormalities, muscle contractures and neurological impairments(10). ZIKV-exposed children have impaired neurodevelopment, but the long-term effects in ZIKV exposed infants remains an active area of investigation as children exposed during the 2015-2016 outbreak in the Americas are now school aged(11–13). Nonhuman (NHP) models of ZIKV infection mimic the many routes of ZIKV infection, mild human disease, and recapitulate vertical transmission and severe aspects of CZS(14–18). In NHP models of ZIKV vertical transmission, fetal loss was reported in 26% of ZIKV exposed animals and suggests fetal loss in asymptomatic ZIKV infected women may go underreported(16). Moreover, altered myelination in normocephalic fetuses following maternal-to-fetal ZIKV transmission argues ZIKV infection *in utero* can impact pre- and post-natal neurologic development(19). Therefore, the NHP is an ideal model for understanding the mechanisms of human ZIKV infection and for testing ZIKV vaccines.

Innate and adaptive immune responses are important for ZIKV viral clearance and protection against re-infection(20, 21). We and others have found that circulating monocytes and dendritic cells are the major cellular blood targets of ZIKV infection in humans and NHP and can contribute to disease pathogenesis by the production of inflammatory mediators(22–24). Yet, these cells are also potently activated during ZIKV infection and contribute to the antiviral type I interferon response(25). Blood monocyte frequencies increase during HIV infection and although antiretroviral therapy (ART) reduces total monocyte frequencies in PLWH, inflammatory monocytes remain elevated(26). We therefore hypothesized that altered innate immune responses during HIV infection could promote susceptibility to ZIKV infection and alter ZIKV pathogenesis. Here, using the pigtail macaque model of HIV/AIDS, we evaluated the impact of simian immunodeficiency virus (SIV) infection on the susceptibility of peripheral blood mononuclear cells (PBMC) to ZIKV infection, ZIKV persistence and host immunity.

## Results

### In vitro ZIKV replication is impaired in PBMC from NHP acutely infected with SIV

Pigtail macaques (n=7) were infected with SIVmac239M and blood was collected prior to and at 2 and 6 weeks post-SIV infection. Both post-infection timepoints are associated with significant declines in peripheral CD4 counts and correspond with SIV peak and viral setpoint, respectively (**Supplemental Figure 1A and C**). To evaluate whether SIV infection alters the permissivity of peripheral blood mononuclear cells (PBMC) to Zika virus (ZIKV) infection, we isolated fresh PBMC from naïve (Pre-SIV) and SIV+ (Weeks 2 and 6) PTM (n=4) and inoculated cells with ZIKV Brazil 2015 (MOI of 2) *ex vivo*. At 4, 24 and 48 hours post-ZIKV infection (hpi), cells and culture supernatants were collected to measure ZIKV RNA and viral titer. Pre-SIV PBMC were permissive to ZIKV infection, as measured by qRT-PCR and plaque assay, with peak viral replication at 24 hpi (**Figure 1A-B, Supplemental Figure 2**). ZIKV RNA levels in Week 2 SIV+ PBMC were significantly decreased at 24 and 48 hpi compared to pre-SIV PBMC, while Week 6 SIV+ PBMC had similar ZIKV RNA levels to that in pre-SIV PBMC (**Figure 1A**). The kinetics of ZIKV replication in pre-SIV PBMC were similar in PBMC derived from SIV^+^ PTM; however, significantly lower levels of infectious virus were observed at 24 and 48 hpi in Week 6 SIV+ PBMC (**Figure 1B, Supplemental Figure 2**). Supernatants from ZIKV-infected PBMC cultures were subjected to multiplex immunoassay to measure cytokine and chemokine concentration changes at 4, 24, and 48 hpi. All cultures accumulated the pro-inflammatory cytokines MCP-1 and VEGF-A during the 48 hr post-infection period; however, there was only a trend for an increase of IL-5 (p = 0.061) in Week 2 SIV-infected cultures relative to pre-SIV at 24 hr (**Supplemental Figure 3**). These data suggest that cells from acutely SIV-infected animals are less permissive to ZIKV infection.

**Figure 1.**
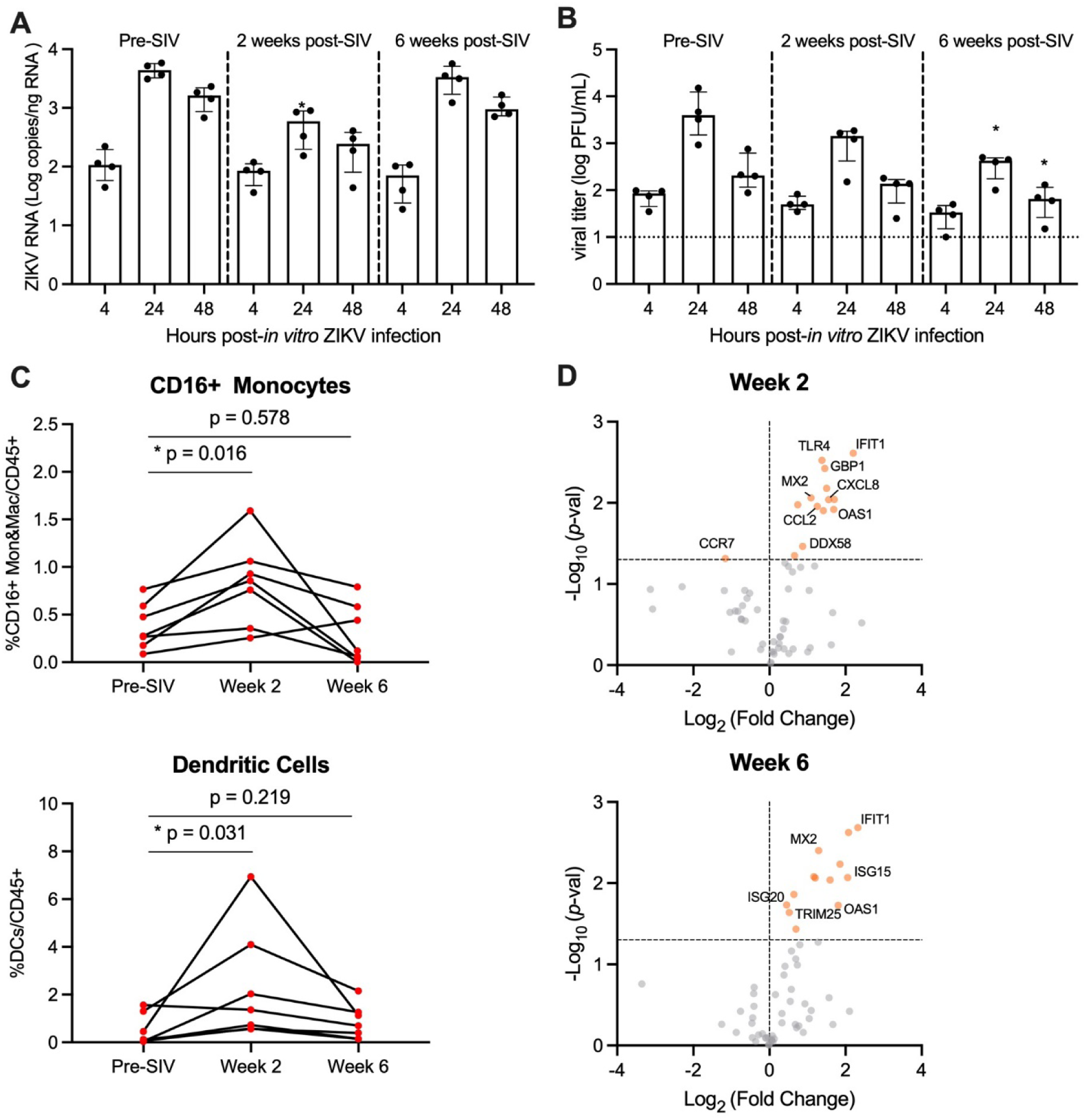
PBMC from SIV-infected PTM are less permissive to *in vitro* ZIKV infection. (**A**-**B**) Peripheral blood mononuclear cells (PBMC) were isolated from pigtail macaques prior to and at 2 and 6 weeks post-SIV infection and infected *in vitro* with ZIKV Brazil at a multiplicity of infection (MOI) of 2. Cells and supernatant were harvested 4, 24, and 48 hours post-infection. (**A**) Quantitative real-time PCR (qRT-PCR) for ZIKV RNA in PBMC. (**B**) Plaque assay for infectious virus. (**A-B**) Medians with interquartile ranges are shown. Kruskal-Wallis test versus pre-SIV levels, p-values * ≤ 0.05. (**C**) Frequency of CD16+CD14+ monocytes and macrophages (top panel) and dendritic cells lower panel) in blood from uninfected and SIV-infected pigtail macaques. Wilcoxon matched-pairs signed rank test, p-values ≤ 0.05 considered significant. (**D**) Gene expression of PBMC in blood at Week 2 post-SIV (top panel) and Week 6 post-SIV (bottom panel). t-test between each time-point relative to Pre-SIV, p-values *<0.01 shown by orange dots.

### Expansion of ZIKV cellular targets in the blood during acute SIV infection

Monocyte frequencies increase in the blood during HIV and SIV infection and are the primary targets of ZIKV infection(22–24, 26). Since ZIKV replication was reduced in PBMC derived from SIV-infected animals, we wanted to determine whether this may be due to decreased ZIKV cellular targets. To test this, we evaluated innate cells in fresh blood from SIV+ and SIV-PTM by flow cytometry. During acute SIV infection, there was a median 2.7-fold increase in the frequency of CD16+ monocytes and a median 9.0-fold increase in the frequency of dendritic cells (DCs) at Week 2, with levels returning to pre-infection levels at Week 6 (**Figure 1C**). AXL, a TAM receptor tyrosine kinase, expressed on monocytes, macrophages, and DCs mediates ZIKV entry into human glial, endothelial, and fetal endothelial cells(34–37). *In vitro* HIV infection of monocyte-derived macrophages causes a 1.5-fold increase in AXL gene expression(38), but it remains unknown if this also occurs *in vivo*. *Ex vivo* AXL expression on CD16+ monocytes and DCs was unchanged at Weeks 2 or 6 post SIV-infection (**Supplemental Figure 4**). Thus, PBMC from SIV-infected PTM have similar/greater levels of ZIKV cellular targets in comparison to naïve PTM and these cells express similar levels of surface AXL, but are less vulnerable to ZIKV infection *ex vivo*. We next hypothesized that anti-viral responses induced early within SIV infection could influence ZIKV permissibility. To test this, we used a targeted custom-built NanoString nCounter gene expression assay to investigate a panel of immune-related genes (84 genes) in PBMC collected at pre-SIV and at Weeks 2 and 6 post-SIV infection. Differential gene expression analysis using a t-test was performed for each time-point post-SIV infection relative to pre-SIV PBMC identified several innate immune and interferon stimulated genes (ISGs) that were significantly upregulated (**Figure 1D, Supplemental Table 4**). With SIV infection, several genes related to innate immunity were upregulated in expression compared to pre-SIV PBMC. Notably, *IFIT1*, *MX2*, and *OAS1* ISGs were increased in expression at both Weeks 2 and 6 post-SIV. *ISG20* and *ISG15* were significantly upregulated in expression at Week 6, while *CXCL8* and *CCL2* encoding monocyte chemoattractant protein were significantly upregulated at Week 2 post-SIV. Retinoic acid-inducible gene-I (RIG-I) signaling activates the expression of these antiviral genes that, in turn, are known to restrict ZIKV replication(39, 40). These data indicate that despite the presence and expansion of ZIKV cellular targets during acute SIV infection, increased innate immune responses in PBMC could render monocytes refractory to ZIKV infection.

### ZIKV co-infection does not significantly impact peripheral SIV disease progression

SIV-infected PTM were co-infected with ZIKV at 63 days (9 weeks) post-SIV infection (SIV+ZIKV+) and compared to SIV-naïve PTM infected with ZIKV (SIV-ZIKV+) (**Figure 2A**). This timepoint post-SIV was selected as an early chronic phase of SIV infection and corresponds with the establishment of viral setpoint (median SIV viremia 5.41 (1.63-6.18) log_10_ copies/mL of plasma) and evidence of immunosuppression including lowered, yet stable peripheral CD4 counts (median 399 (333–901) cells/μL of blood) and decreased frequencies of CD4 T-cells in the gut mucosa relative to SIV-naïve controls (**Supplemental Figure 1C-E, Supplemental Table 1**).

**Figure 2.**
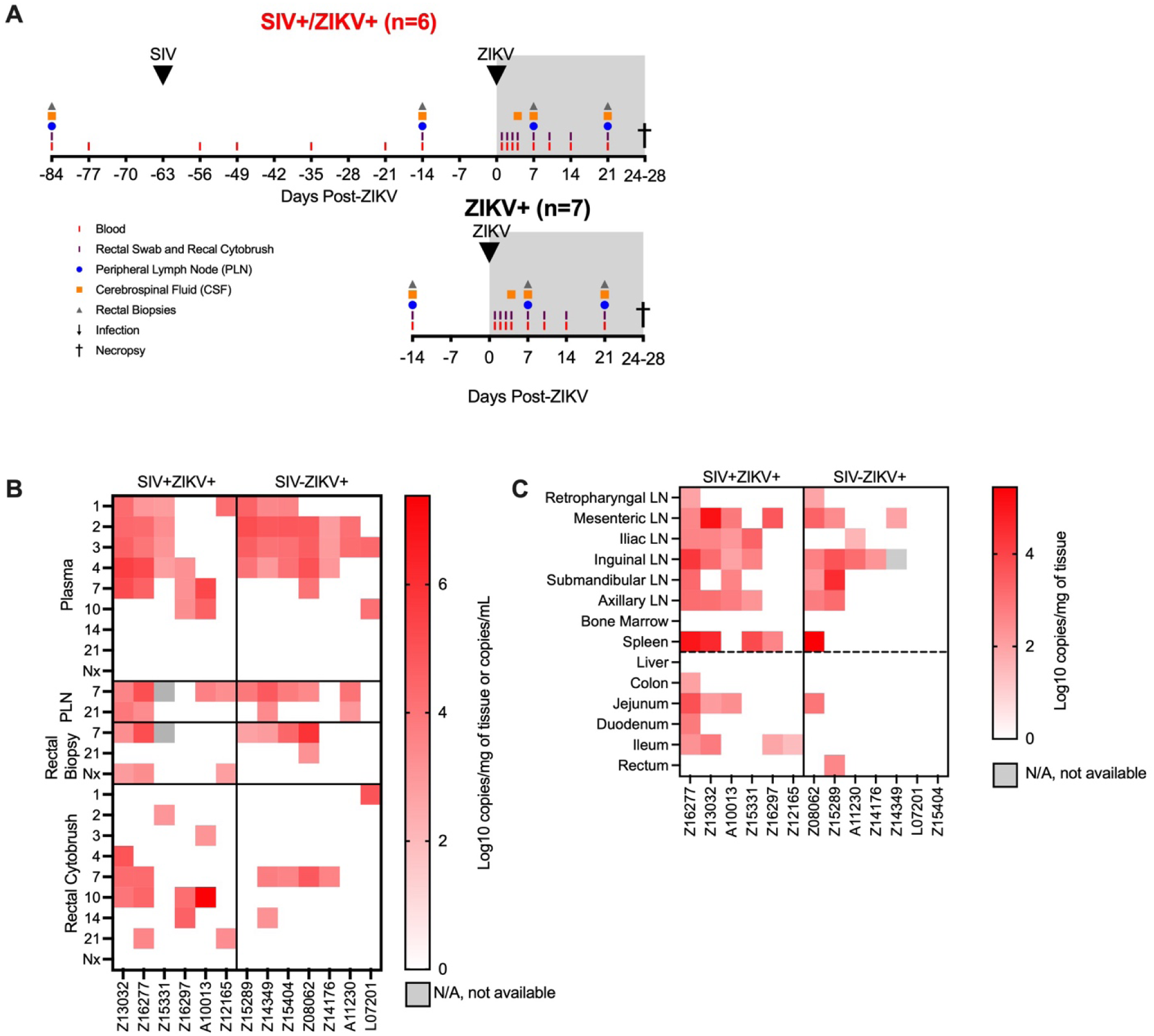
ZIKV viremia is delayed and protracted and ZIKV viral burden more persistent in gastrointestinal tissues in SIV-infected macaques. (**A**) Study design of longitudinal blood and tissue sampling following SIV and ZIKV infections in pigtail macaques. Initially n=7/group were infected with ZIKV, however 1 animal in the SIV+ group displayed no evidence of ZIKV replication and thus was excluded from all post-ZIKV analysis. (**B**) Quantitative real-time PCR (qRT-PCR) for ZIKV RNA in longitudinal samples from plasma, peripheral lymph node (PLN), rectal biopsies, and rectal cytobrush until necropsy (Nx). Virus was not detected in any longitudinal cerebrospinal fluid (CSF). (**C**) ZIKV RNA in tissues collected at necropsy 24-28 DPI. Virus was not detected in brain tissue (brainstem, hippocampus, frontal lobe, parietal lobe, and occipital lobe) of any animal.

Blood, peripheral lymph nodes (PLN), cerebrospinal fluid (CSF), and rectal samples (biopsy, cytobrush) were longitudinally collected according to **Figure 2A** for 4 weeks until the time of necropsy. In the SIV+ZIKV+ cohort, SIV viremia and peripheral CD4 counts remained stable post-ZIKV coinfection and there was no evidence of enhanced gut barrier dysfunction (**Supplemental Figure 1C-E, 5A**). The most notable histologic findings were in Z16197, who had the syndrome proliferative-occlusive pulmonary arteriopathy with thrombosis and infarction, which is a retroviral-strain-associated disease that was likely secondary to the SIV infection **(Supplemental Table 2)**. Overall, these findings suggest that acute ZIKV co-infection does not have a significant effect on SIV viral replication or disease progression.

### SIV infection promotes delayed ZIKV viremia and increases ZIKV persistence in the gut mucosa

To assess the impact of the early chronic phase of SIV infection on ZIKV pathogenesis and tissue tropism, ZIKV burden was evaluated in longitudinal specimens and in tissues at necropsy by qRT-PCR. ZIKV RNA was not detected in one animal in the SIV+ZIKV+ cohort (Z14109) in any longitudinal sample tested nor in any necropsy tissue (**Supplemental Table 7-8**); therefore, there was no evidence of productive ZIKV infection, and the animal was excluded from all subsequent post-ZIKV analysis. In the SIV-ZIKV+ cohort, plasma viremia peaked 2-4 days post infection (dpi) (median 3 dpi) and ZIKV was cleared in the plasma in most animals (6/7) by 7 dpi (**Figure 2B**), and the mean viral kinetics in this cohort was consistent with our previous findings in ZIKV-infected PTM(24). In contrast, in SIV+ZIKV+ PTM, peak ZIKV viremia was variable (1-10 dpi; median 4 dpi) and virus was still present in most animals (4/7) by 7 dpi (**Figure 2B**). Accordingly, median plasma viral loads at 2 and 3 dpi trended to be higher or significantly higher (2 dpi, p = 0.108; 3 dpi p = 0.050) in the naïve animals compared to SIV+ZIKV+ PTM and, but instead trended higher (p=0.054) in SIV+ZIKV+ PTM at 7 dpi (**Supplemental Figure 6A**). Together, these data indicate that initial ZIKV viremia and clearance in the periphery are delayed in SIV-infected macaques.

We next examined ZIKV burden in longitudinal PLN, rectal tissue biopsies, and CSF collected at days 4, 7 and 21 post-ZIKV challenge, as these are known sites of ZIKV tropism(14, 24). In PLN, ZIKV RNA was detected in most animals (SIV-5/7; SIV+ 4/6) and in a proportion of animals (SIV-2/7; SIV+ 2/6) at 7 and 21 dpi, respectively (**Figure 2B**). In rectal biopsy tissue, ZIKV RNA was detected in 4/7 of SIV- and 2/6 of SIV+ animals at 7 dpi, and at the time of necropsy (24-18 DPI) was detected in 1/7 of SIV- and 3/6 of SIV+ animals at necropsy (**Figure 2B, Supplemental Figure 6**), providing evidence for ZIKV persistence in the rectum during SIV infection. In rectal cytobrushes, ZIKV RNA was predominantly detected in SIV-animals at 7 dpi (4/7 PTM) and was sporadically, but more consistently detected in SIV+ animals at 10 dpi (4/6 PTM) and the total viral burden trended (p = 0.171) to be higher during SIV+ZIKV+ co-infection (**Figure 2B, Supplemental Figure 6**). ZIKV RNA was not detected in CSF of any animal at any timepoint throughout the study (**Supplemental Table 7**). Overall, these data suggest that ZIKV infectivity of the gut mucosa may persist during untreated SIV infection.

To evaluate the impact of SIV infection on ZIKV tropism and persistence in tissues, we measured ZIKV viral burden in lymphoid, gut mucosal, and neuronal tissues at necropsy (24-28 dpi). In both cohorts, ZIKV RNA was detected in both lymphoid and gastrointestinal tissues, but not in brain tissue (brainstem, hippocampus, frontal lobe, parietal lobe, and occipital lobe), which further corroborates findings in the CSF suggesting that ZIKV was not neurotropic in the animals (**Figure 2C, Supplemental Table 8**). The total number of ZIKV+ tissues at necropsy trended higher in SIV+ZIKV+ animals (p = 0.108) compared to SIV-ZIKV+ animals (**Supplemental Figure 7A**). Upon further examination, ZIKV RNA was detected in at least one lymphoid tissue in most animals from each cohort (5/6 SIV+ZIKV+; 5/7 SIV-ZIKV+) at necropsy and the median number of positive tissues at necropsy was 4.5 for SIV+ and 1.0 for SIV-PTM (**Figure 2C, Supplemental Figure 7**). There was no significant difference in the number of positive lymphoid tissues or in the individual or total ZIKV viral burden within lymphoid tissues (**Supplemental Figure 7**). At necropsy, ZIKV RNA was detected in at least one gut associated tissue in a majority (5/6) of SIV+PTM but only in 2/7 SIV-PTM (**Figure 2C**). In accordance, the median number of ZIKV+ gastrointestinal tissues trended (p = 0.057) to being higher in SIV-infected PTM (**Supplemental Figure 7**) and there trended to be a greater total (p = 0.125) gut ZIKV viral burden during SIV co-infection (**Supplemental Figure 7**). Collectively, these data suggest that ZIKV distribution and viral burden is similar in the lymphoid tissues, but that SIV infection could contribute to greater ZIKV persistence in gut mucosal tissues.

### Delayed and dampened expansion of ZIKV cellular targets in blood corresponds with the recruitment of cellular targets to tissues during SIV-ZIKV coinfection

As there was evidence for altered ZIKV pathogenesis in SIV-infected animals, we next wanted to evaluate whether this was associated with changes to ZIKV cellular targets or the immune response. Humoral responses are important for the control of ZIKV infection(41) and we measured anti-ZIKV envelope IgG responses in longitudinal plasma samples. SIV-ZIKV+ animals generated robust binding IgG antibodies that were detected at 7 dpi and peaked at 14 dpi. In contrast, anti-ZIKV envelope IgG responses were lower overall in SIV+ZIKV+ animals (AUC, p = 0.014) (**Figure 3A**). Despite the difference in anti-ZIKV IgG between the two cohorts, there was a similar level of neutralizing antibodies (NAb) generated against ZIKV at necropsy (7/7 SIV-ZIKV+, 6/7 SIV+ZIKV+), with no significant difference in the overall NAb level between groups (**Figure 3B**).

**Figure 3.**
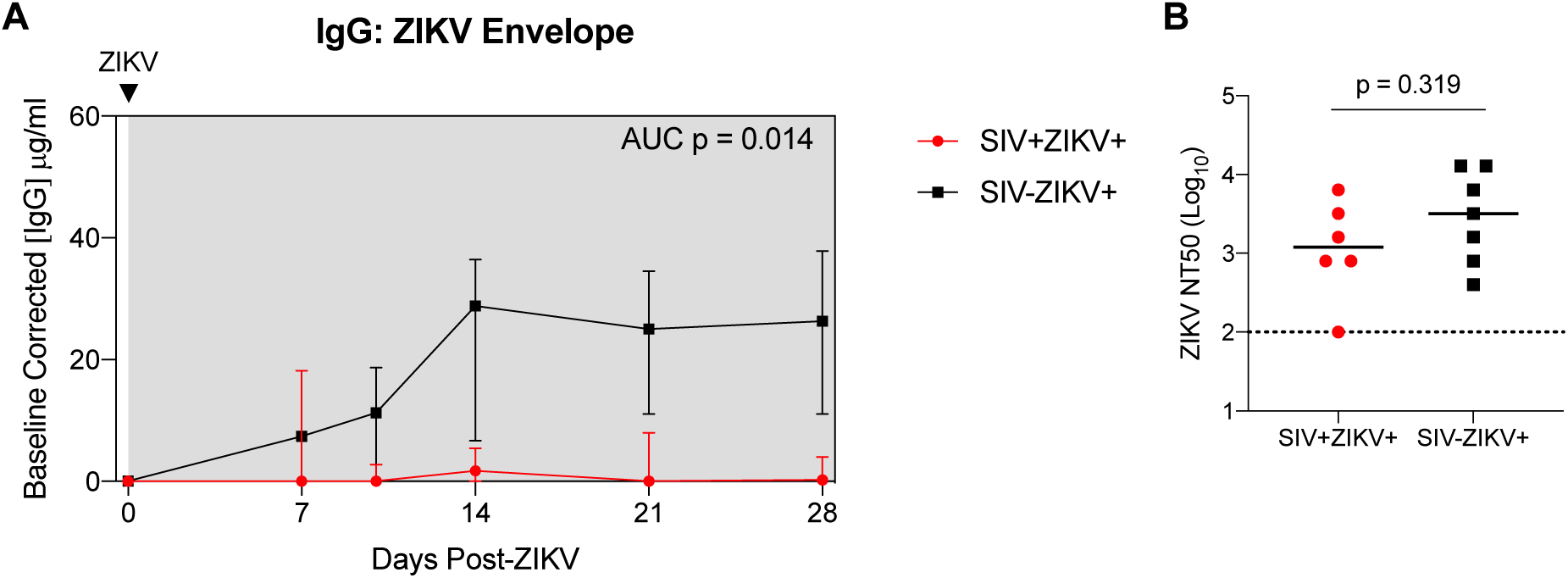
SIV infection may impair anti-ZIKV immunity. (**A**) Longitudinal plasma concentrations of anti-ZIKV envelope IgG as determined by ELISA. AUCs were calculated from day 10 to 28. Medians with interquartile ranges are displayed. (**B**) Zika virus neutralization antibody titers (NT50 values) evaluated at necropsy. The line represents the median and the dotted line represents is the limit of detection. (**A-B**) Mann-Whitney test comparison between groups.

Consistent with our previous findings(24), there is rapid and robust expansion of inflammatory CD16+ (non-classical and intermediate) and CD16-(classical) monocytes in the blood in the first few days after ZIKV infection (median peak 2 dpi), which corresponds to peak ZIKV viremia (**Figure 4A, Supplemental Figure 8**). In contrast, the expansion of CD16+ and CD16-monocytes was severely dampened and delayed during SIV infection (median peak 8.5 days) (**Figure 4A, Supplemental Figure 8**) and corresponded with the delayed peak ZIKV viremia observed in these animals (**Figure 2B**). Cellular analysis in the tissues revealed that CD16-monocytes/macrophages were robustly and significantly recruited to the rectum and PLN in SIV-negative animals, whereas in contrast CD16+ monocytes/macrophages were recruited to the tissues during SIV+ZIKV+ co-infection (**Figure 4A, Supplemental Figure 8**). Additionally, AXL expression was not significantly changed on ZIKV cellular targets in whole blood, rectum and PLN in either group post-ZIKV infection (**Supplemental Figure 9**). These findings suggest that increased recruitment of inflammatory monocytes and macrophages to lymphoid and gastrointestinal tissues during SIV infection, which are also cellular targets of Zika virus infection, may contribute to ZIKV viral persistence at these sites.

**Figure 4.**
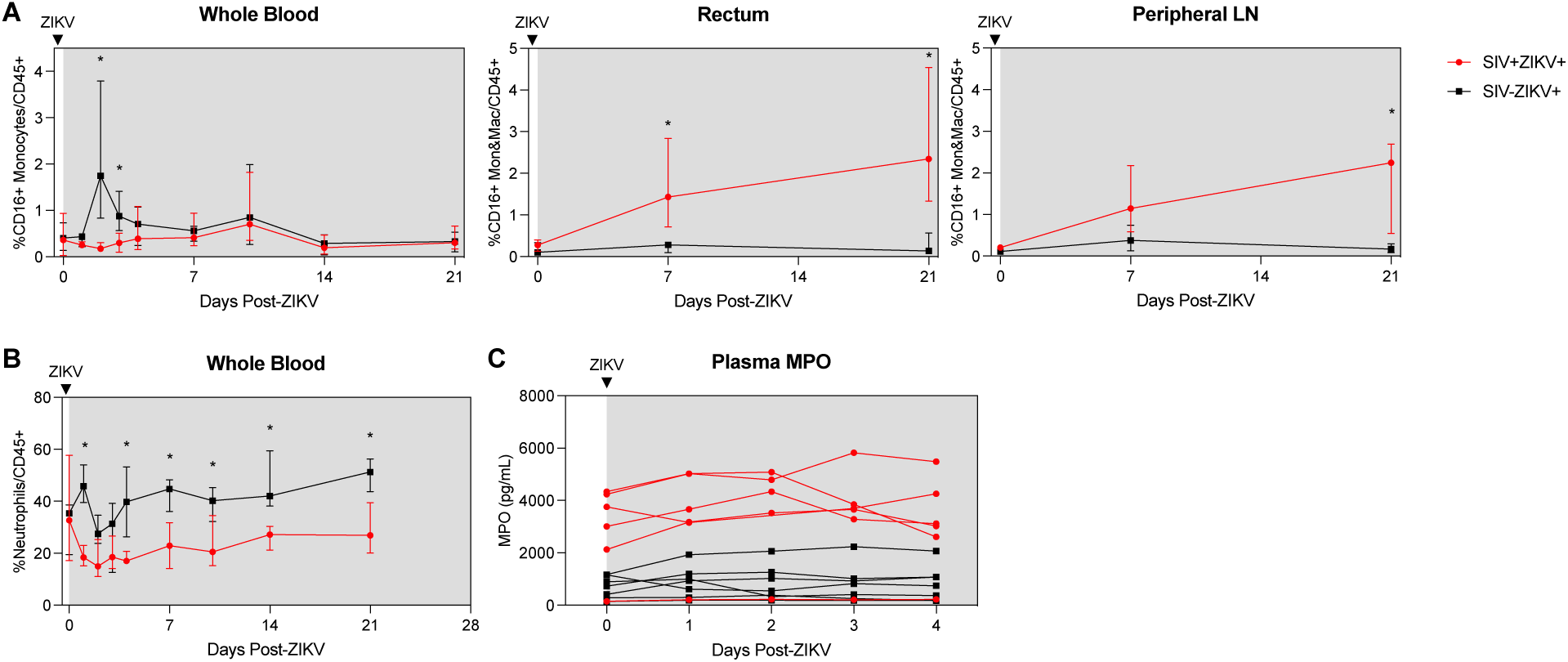
Post-ZIKV recruitment of CD16+ monocytes and macrophages is dampened in the periphery, but enhanced in tissues in SIV-infected macaques. (**A**) Frequency of CD16+CD14+ monocytes and macrophages in blood (left panel), rectum (center panel), and peripheral lymph node (right panel) after ZIKV infection. (**B**) Frequency of neutrophils in blood after ZIKV infection. (**C**) Concentration of myeloperoxidase (MPO) in plasma as measured by ELISA. (**A-C**) Medians with interquartile ranges are shown. Mann-Whitney test between group, p-values * ≤ 0.05.

Neutrophils are important for ZIKV dissemination and pathogenesis(24, 42, 43). Blood neutrophils declined during SIV+ZIKV+ co-infection starting at 1 dpi and were significantly lower in frequency in comparison to SIV-ZIKV+ animals through 21 dpi (**Figure 4B**). The frequencies of neutrophils in the tissues were also lower in SIV+ZIKV+ compared to SIV-ZIKV+ animals in the rectum at 21 dpi, with no differences in neutrophil frequencies observed within the PLN (**Supplemental Figure 10**). To assess neutrophil function, we evaluated plasma concentrations of myeloperoxidase (MPO), a neutrophil granule and marker of inflammation, during the first 4 days of ZIKV infection. Interestingly, prior to ZIKV infection, elevated levels of MPO were detected in 5/6 SIV+ animals while lower concentrations of MPO were detected in 7/7 SIV-animals (**Figure 4C**). Post-ZIKV infection, concentrations of plasma MPO continue to be significantly higher in SIV+ZIKV+ animals in comparison to SIV-ZIKV+ animals (**Figure 4C**). These data suggest that neutrophils are more inflammatory during SIV infection and there is an impairment of neutrophil recruitment to the blood and impaired trafficking to gastrointestinal tissues post-ZIKV.

### SIV-ZIKV co-infection induces chronic innate immune activation

The inflammatory response to ZIKV in the plasma was evaluated using a multiplex immunoassay. In both groups, ZIKV induced pro-inflammatory responses, characterized by transient increases in interleukin-1 receptor agonist (IL-1RA), monocyte chemoattractant protein-1 (MCP-1), and vascular endothelial growth factor A (VEGF-A) (**Supplemental Figure 11**), with no major differences between groups. Although we found no evidence of ZIKV infection in neuronal tissue, neuroinflammation was evaluated in longitudinal CSF specimens. CSF concentrations of sCD14, a marker of neuroinflammation, did not change with ZIKV infection in either group (**Supplemental Figure 5B**). IL-1RA, IL-6, IL-8, and VEGF-A were elevated in several animals across groups at varying timepoints (**Supplemental Figure 11**). Transient increases in IL-8 were detected in the CSF of a few SIV-ZIKV+ animals after ZIKV infection, which is evidence of neuroinflammation. At necropsy, two animals in the ZIKV+ group (L07201 and Z08062) had mild, multifocal demyelination and axonal loss in the brainstem, which may be a result of neuroinflammation despite no evidence of ZIKV infection at this site (**Supplemental Table 2, 8**). Overall, no significant differences between SIV+ vs SIV-groups were found for any analytes detected in plasma or CSF during ZIKV infection, suggestive that SIV infection does not enhance ZIKV-induced systemic inflammation or neuroinflammation.

To further examine SIV effects on immune responses to ZIKV, we performed targeted gene expression analysis on longitudinal PBMC specimens collected post-ZIKV infection using a custom-built NanoString Code Set for interrogating 64 genes marking innate activation, inflammatory and interferon (IFN) responses. Changes in gene expression were determined at each time-point post challenge in comparison to uninfected timepoints. Gene expression was not significantly changed 7 weeks post-SIV infection relative to naïve PBMC. This indicated that while SIV infection stimulated gene expression, the changes just prior to ZIKV challenge were not significantly different from baseline levels. Following ZIKV challenge, a total of 23 genes were significantly differentially expressed (22 upregulated and 1 downregulated), 14 genes in SIV-ZIKV+ (13 up and 1 down), 19 genes in SIV+ZIKV+ (19 up), and 10 genes in both groups (10 up) **(Figure 5B, Supplemental Table 5).** ZIKV induced robust innate immune gene activation in SIV-ZIKV+ animals during acute infection (2-4 dpi), with the expression of most genes returning to baseline by 7 dpi (**Figure 5A**). The kinetics of the gene expression mirrors peak ZIKV viremia (median 3 dpi) and time to viral clearance (median 7 dpi). While these innate immune genes were also strongly upregulated in SIV+ZIKV+ co-infected animals during acute infection (2-4 dpi), in contrast to the SIV-ZIKV+ group, the gene signature was maintained throughout infection and remained highly expressed 7-21 dpi. The kinetics of gene expression during SIV+ZIKV+ co-infection corresponds to the shift in peak ZIKV viremia (median 4 dpi) and viral clearance (median 10 dpi). We also found that the expression of 10 genes were significantly different between SIV+ZIKV+ and SIV-ZIKV+ animals, primarily at 14 and 21 dpi, and predominantly consisted of genes associated with type I IFN signaling and ZIKV viral control (*ISG15, IFIT1, MX1, ISG20, IRF7)* (**Figure 5C, Supplemental Table 6**). These data demonstrate that SIV-ZIKV co-infection leads to persistent upregulation of genes associated with inflammation and innate immune activation in whole blood. This chronic hyperactivated innate immune state in response to ZIKV co-infection is likely a contributing factor in the impaired peripheral ZIKV clearance and ZIKV persistence in tissues.

**Figure 5:**
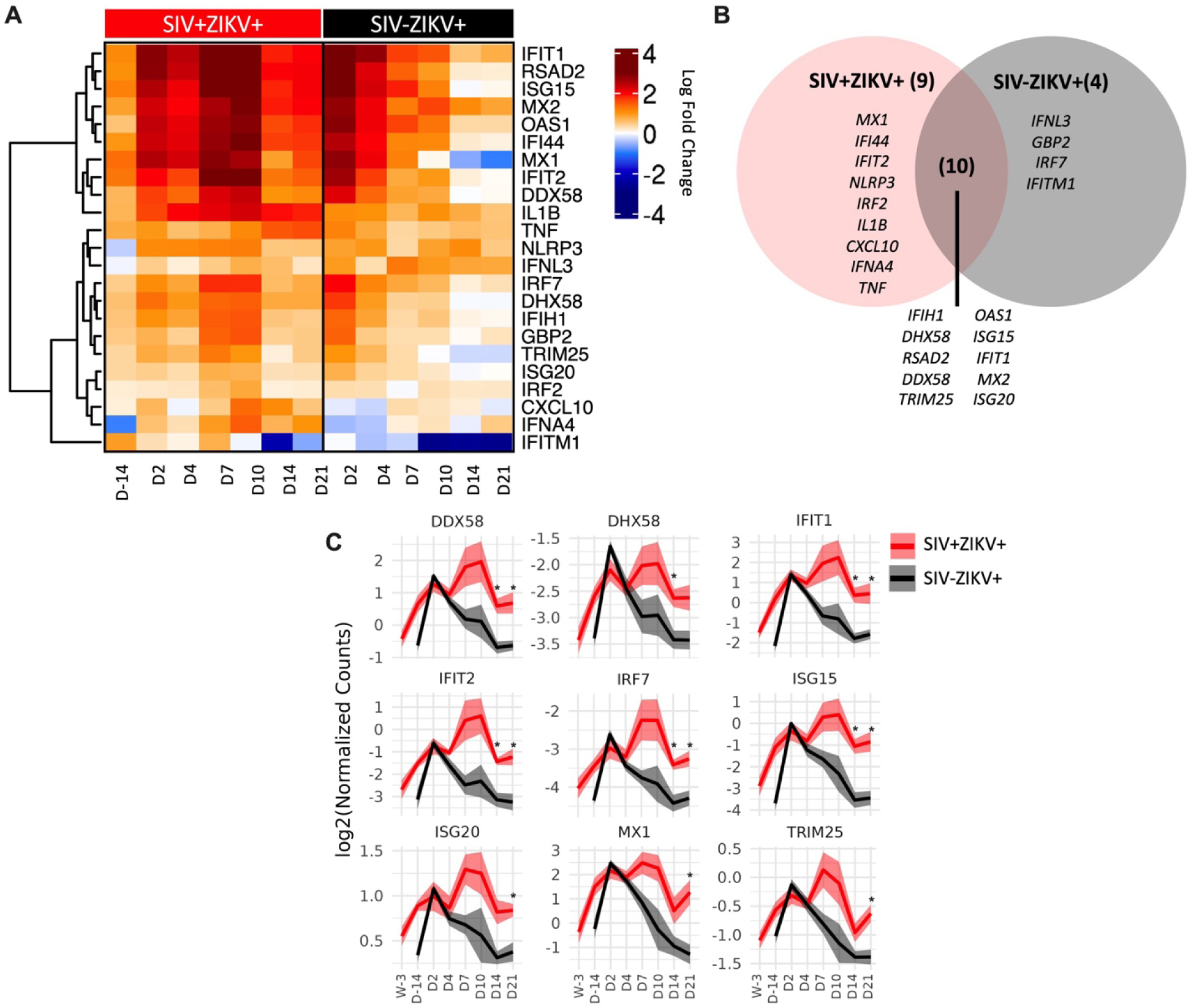
SIV-ZIKV co-infection induces chronic innate immune activation in PBMC. (**A**) Heatmap showing the LFC expression of 23 genes that were significantly different (*p*-value<0.01) in at least one time point and one group. LFC expression for the SIV+ZIKV+ group is relative to pre-SIV (Wk-3). LCF expression for the SIV-ZIKV+ group is relative to pre-ZIKV (D-14). Genes’ LFCs were clustered using Pearson and Ward.D2. (**B**) Venn diagram of shared and unique genes that were significantly upregulated post-ZIKV. (**C**) Line plots of select gene kinetics representing the mean of all SIV+ZIKV+ (red) or SIV-ZIKV+ (black) animals. The log2 normalized counts are plotted at each time point. The line represents the mean and the standard error is shown as the confidence interval around the mean. p-values *<0.01 indicates a significant difference between SIV+ZIKV+ and or SIV-ZIKV+ at a specified time point.

## Discussion

In this study, we aimed to determine whether HIV-induced immunosuppression impacts ZIKV pathogenesis and investigated this using *in vitro* and *in vivo* models of SIV-ZIKV co-infection. We identified that peripheral ZIKV cellular targets, including CD16+ monocytes, increase during acute infection but contract to pre-infection levels during early chronic stages of SIV infection (**Figure 6**). Despite AXL being an important cellular receptor of ZIKV infection, we found no change in AXL expression on monocytes in whole blood or tissues of SIV+ animals following ZIKV challenge. This indicates that enhanced ZIKV persistence during SIV infection is unlikely to be caused by increased receptor engagement on cellular targets. Interestingly, PBMC from acutely SIV infected NHP exhibit an anti-viral gene expression profile that renders the cells refractory to ZIKV co-infection *in vitro.* We also demonstrated *in vivo* that SIV infection modulates the innate and adaptive immune response to ZIKV co-infection and creates a hyper inflammatory state that could contribute to prolonged viremia and impaired viral clearance from the tissues, particularly in the gastrointestinal tract (**Figure 6**). This study further implies that PLWH or other immunocompromised individuals co-infected with ZIKV have the potential for longer ZIKV transmission periods.

**Figure 6:**
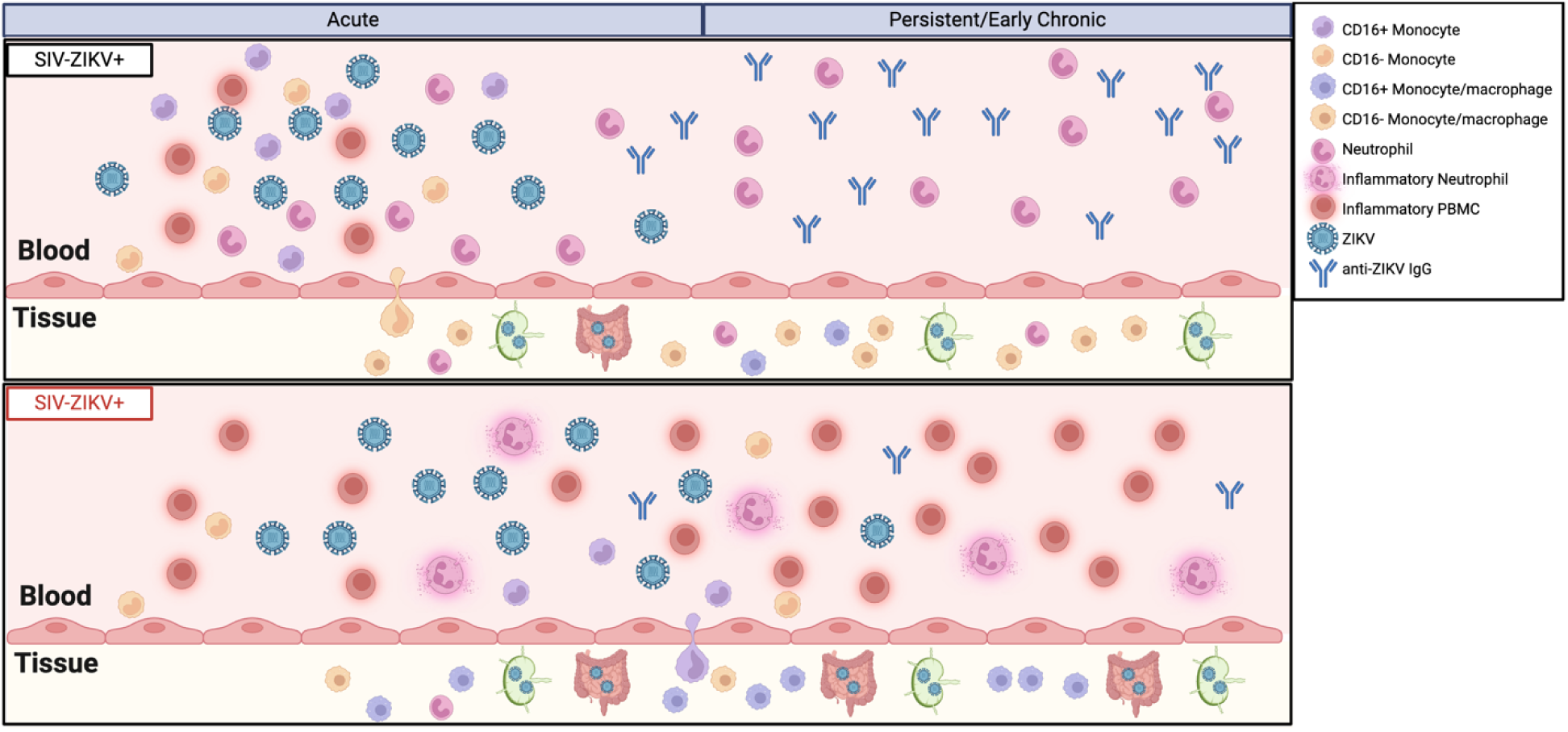
Model of innate immune impairment and ZIKV persistence during SIV infection. The model depicts impaired induction of monocyte recruitment, presence of inflammatory neutrophils, and chronic immune activation in the periphery. These findings, coupled with the absence of anti-ZIKV IgG, suggest a mechanism for delayed clearance of ZIKV viremia. ZIKV persistence in gastrointestinal tissues is enhanced in an immunocompromised state and the preferential recruitment of CD16+ monocytes/macrophages into tissue may be a contributing mechanism. Created with biorender.com.

The clinical impact of HIV co-infection on flavivirus infection remains unclear. For example, some studies report less, while others report more severe disease during DENV-HIV co-infection(44–46). ZIKV infection during pregnancy in women living with HIV is of great concern: one study reported 12% of ZIKV exposed infants to women living with HIV had CNS abnormalities(47), as compared to reports of 5-8% in ZIKV exposed infants from HIV^-^ women(48), indicating there may be greater risks of neuropathologies in HIV-ZIKV exposed infants. These flavivirus co-infection studies in PLWH are limited by low patient numbers, inconsistent incorporation of ART and HIV disease status, and parameters are primarily restricted to measurements in the blood. Studies examining ZIKV infection in SIV or SHIV infected NHP have helped fill this clinical gap. One study found that pregnant female macaques with treated SIV infection (SIVmac239), had similar rates of pregnancy loss due to ZIKV exposure (ZIKV-DAK) when compared to SIV naïve animals(49). Whether rates of fetal loss are similar in those with untreated SIV infection remains an area of potential future investigation. Two studies have investigated ZIKV co-infection in non-pregnant rhesus macaques with untreated SIV/SHIV infection(50, 51). Bidokhti et al., reported no differences in ZIKV virema with SIV/SHIV infection; however, interpretations from this study are limited by low animal numbers and a lack of contemporaneous controls(51). Notably and similar to our findings here, Vinton et al., observed delayed peak ZIKV viremia and clearance in SIV-infected rhesus(50). Higher levels of ZIKV RNA were detected in lymph nodes from SIV+ rhesus at 27 dpi(50), and while we detected persistence of ZIKV in SIV+ and SIV-pigtail macaque, we found no significant differences in viral burden between groups. These differences may be due, in part, to the sensitivities of the methods used, fluorescence in situ hybridization in Vinton et al., versus qRT-PCR in our study. Notably, our study uniquely provides additional insight that SIV infection promotes ZIKV persistence in gastrointestinal tissues, a tissue compartment that was not reported in previous studies. Major differences in experimental design between previous reports and our study here include 1) different species (rhesus versus pigtail macaques), 2) the time of ZIKV co-infection (3 weeks or 6-7 months post-SIV versus 9 weeks post-SIV), and 3) strain of ZIKV (Nicaragua/2016 and PRVABC59 versus Brazil_2015). Collectively, our study and that of Vinton et. al suggest that untreated acute SIV infection alters ZIKV viral kinetics and enhances ZIKV persistence or viral burden in tissues. This suggests that immunocompromised individuals may have longer ZIKV transmission periods or altered disease pathogenesis and should be primary candidates for ZIKV vaccines.

ZIKV and West Nile Virus (WNV) can invade the central nervous system, yet the mechanisms of neuropathogenesis are not well understood(52, 53). ZIKV RNA in the CSF or brain is detected infrequently but consistently reported in rhesus and cynomolgous macaques(20, 25, 54–56). In contrast, ZIKV RNA in the CSF or CNS was not detected in adult pigtail macaques reported here and in our previous report(24). This suggests that the frequency or severity of ZIKV neuroinvasion may vary between macaque species and that different host factors could play a role. Evidence for neuroinflammation is reported in ZIKV infected NHP, even when ZIKV RNA is undetectable in the CSF(54, 57–59). Increases in inflammatory infiltrates into the CSF of ZIKV infected rhesus, including IL-15, MCP-1, G-CSF, and CXCL12, are previously reported(57, 58). Here, we also observed that IL-1RA, IL-6, and IL-8 concentrations were elevated in the CSF; however, these responses were highly variable between animals. It is plausible that the differences in observed cytokine milieus between our study in pigtail macaques and those in rhesus macaques are due to assay sensitivities and cross-reactivity levels across assays. However, this could point to potential differences in neuroinflammatory responses between macaque species. In PLWH, increased WNV neuroinvasion occurs(60–62) and supports that HIV-ZIKV co-infection could similarly result in increased targeting of ZIKV to the CNS, resulting in higher rates of neurological pathologies. Our studies here provide no evidence that SIV infection promotes increased ZIKV neuroinvasion or neuroinflammation, however additional studies in other NHP species are needed to fully assess this risk.

Many promising ZIKV vaccine candidates in the pre-clinical pipeline rely on the induction of antibodies and T-cells to mediate protection(63). Studies in NHP revealed that CD8 T-cells are not required for protection against primary or secondary ZIKV infections(25, 64). Another study in NHP further showed that the impaired generation of anti-flavivirus humoral responses in CD4 depleted macaques infected with DENV and ZIKV, shapes the quality of responses to a tertiary flavivirus exposure(65). Here, our study is in congruence with Vinton et al.(50), and demonstrates that the generation of anti-ZIKV adaptive immunity is impaired in SIV infected animals that exhibit CD4 immunodeficiency. Further studies are needed to elucidate whether the immune responses generated during CD4 immunosuppression are sufficient for protection against homologous and/or heterologous re-exposure to ZIKV. This will improve our ability to identify groups at risk for re-infection and to create effective vaccines in various states of immunosuppression, including those for the elderly, pregnant women, and immunocompromised individuals.

The innate immune response is the primary defense against flavivirus infection, but if dysregulated or uncontrolled, it can lead to enhanced viral pathogenesis. This dichotomous role is further complicated during flavivirus infection, as innate immune cells, including monocytes and dendritic cells, are targets of ZIKV, WNV, and DENV infection. In our *in vitro* co-infection model, we found that PBMC from SIV infected animals were less permissive to ZIKV infection, and in our *in vivo* model, ZIKV replication was delayed in the periphery. Initially cells from immunosuppressed individuals may be less permissive to ZIKV infection due to an antiviral state and this may contribute to a slower establishment of ZIKV infection. However, once established, the ability to recruit immune cells critical for clearing the ZIKV infection is impaired and dysregulated during immunosuppression, promoting dissemination and persistence of ZIKV in the tissues, most notably the gastrointestinal tract (**Figure 6**). Previously, we reported that the infiltration of neutrophils and CD16-classical monocytes/macrophages into tissues may be important for controlling Zika virus replication(24). Here, we further corroborate this finding and demonstrate that CD16+ inflammatory monocytes/macrophages, but not CD16-monocytes/macrophages nor neutrophils, traffick into the tissues of SIV+ZIKV+ animals and correspond to ZIKV persistence (**Figure 6**). Prolonged innate immune activation is a hallmark of chronic infections, including HIV, but can also be observed following acute viral infections. Notably, SARS-CoV-2 infection is predominantly an acute infection, yet in some individuals SARS-CoV-2 virus can persist in the tissues and/or lead to post-acute sequelae of COVID-19 (PASC)(66). PASC is associated with persistent immune activation and PLWH are at greater risk for PASC(67). This suggests that chronic innate immune activation can be a major driver in promoting viral persistence. Our study supports the possibility that chronic innate immune activation could similarly contribute to ZIKV persistence. Due to the variability of immune responses and disease outcomes in our study, additional studies are needed to elucidate the immune mechanisms contributing to flavivirus persistence in states of immunosuppression.

## Materials & Methods

### Study Design and Animal Welfare

A total of 14 male and female pigtail macaques (aged 4-11 years, 6-13 kg) were used. **Supplemental Table 1** details animal characteristics, including MHC haplotypes and experimental vaccination history. Prior to enrollment, all animals were pre-screened and seronegative for the presence of antibodies to West Nile, dengue, and Zika viruses. At least 2 months prior to enrollment into the study, eight animals were previously enrolled in studies in which they received an experimental hepatitis B virus (HBV) vaccine consisting of a combination of CD180 targeted DNA and recombinant protein vaccines comprised of HBV core and surface antigens(27) and/or a replicating RNA COVID-19 vaccine(28) (**Supplemental Table 1**) and were evenly distributed between the control and experimental groups. Seven pigtail macaques were infected intravenously with 10,000 infectious units (I.U.) of SIVmac239M(29) (gift from Dr. Brandon Keele, AIDS and Cancer Virus Program, Frederick National Laboratory for Cancer Research) and then co-infected with ZIKV at 9 weeks. All animals were subject to a Simian AIDS monitoring protocol as defined by the WaNPRC guidelines(30). All animals were inoculated with 5 x 10^5^ PFU of the Brazil_2015_MG strain of ZIKV (GenBank: KX811222.1), as previously described(24). ZIKV RNA was not detected in any specimen tested at any timepoint in one animal in the SIV-infected group (Z14109); therefore, this animal was excluded from all post-ZIKV analysis. All animals were euthanized at the study endpoint at 4 weeks post-ZIKV infection under deep anesthesia, in accordance with the 2007 American Veterinary Medical Association Guidelines on Euthanasia, by administration of Euthasol^®^ (Virbac Corp., Houston, TX). As previously described(24), all animals were housed at the Washington National Primate Research Center (WaNPRC), an accredited facility the American Association for the Accreditation of Laboratory Animal Care International (AAALAC). All animal procedures were approved by the University of Washington’s Institutional Animal Care and Use Committee (IACUC) and were collected and processed as previously described(24) and according to the schematic in **Figure 2A**. Animals were observed daily and full physical exams were conducted at each experimental timepoint, as previously described(24).

### Simian AIDS Measurements

SIV plasma viremia was evaluated by quantitative real time reverse transcription polymerase chain reaction (RT-PCR) by the Virology and Immunology Core at the WaNPRC, as previously described(30), and by the NIAID DAIDS Nonhuman Primate Core Virology Laboratory (NHPCVL) for AIDS Vaccine Research and Development Contract. Complete blood counts (CBC) and serum chemistries were performed by the Research Testing Service (RTS) at the University of Washington Department of Laboratory Medicine. Peripheral CD4 counts were determined from CBC using flow cytometry-based methods by the Virology and Immunology Core at the WaNPRC, as previously described(31).

### Cell culture and virus stock

Peripheral blood mononuclear cells (PBMC) were isolated from NHP whole blood collected pre-SIV inoculation (Wk-3) and at weeks 2 and 6 post-SIV inoculation, as previously described(24). PBMC were maintained in RPMI medium supplemented with 10% fetal bovine serum (FBS; HyClone), 2 mM L-glutamine, 5 mM sodium pyruvate, 1x Antibiotic Antimycotic Solution, and 10 mM HEPES (cRPMI; complete RPMI). RPMI medium used for the ZIKV inoculation was supplemented with 1% FBS, 2 mM L-glutamine, 5 mM sodium pyruvate, 1x Antibiotic Antimycotic Solution, and 10 mM HEPES (iRPMI; infection RPMI). Vero cells (WHO, Geneva, Switzerland) were cultured in complete Dulbecco’s modified Eagle medium (cDMEM) supplemented with 10% FBS, 2 mM L-glutamine, 5 mM sodium pyruvate, 1x Antibiotic Antimycotic Solution, 10 mM HEPES and 1X non-essential amino acids. Vero cells tested negative for mycoplasma contamination. All cells were maintained in a 37°C incubator with 5% CO_2_. Brazil Zika virus stock (GenBank: KX811222.1) was used for the PBMC inoculation.

### ZIKV infection of PBMC

Following overnight incubation, PBMC cell suspensions were prepared, and the cell concentration and viability measured using the Countess 3 Automated Cell Counter (ThermoFisher Scientific). Approximately 6 x 10^6^ PBMC were inoculated with ZIKV at an MOI of 2 in a total volume of 200 µL RPMI infection medium (iRPMI) at 37°C for 2 hours (h). Cells were gently mixed by pipetting at 20 minute (min) intervals during incubation. After 2 h, the cells were spun at 300 relative centrifugal force (rcf) for 3 min at RT and the inoculum carefully removed without disturbing the cell pellet. The cells were washed with 300 µl iRPMI and then resuspended in pre-warmed complete RPMI (cRPMI). A total of 5 x 10^5^ PBMC were added to each well of a 24-well plate containing 1 mL of cRPMI. The plates were returned to 37°C and incubated until the designated time-point for sample collection. At 4, 24, and 48 h post-ZIKV inoculation, supernatants were collected and spun at 300 rcf for 3 min at 4°C. 100 µL Versene solution (ThermoFisher Scientific) was added to each well to dislodge adherent cells from the TC plate. The clarified supernatant was transferred to a new tube and banked at -80°C until further processing by plaque assay or qRT-PCR assay. The Versene solution containing PBMC was added to the PMBC cell pellet from the supernatant and then spun at 300 rcf for 3 min at 4°C. The supernatant was carefully removed from the cell pellet and discarded. The cell pellet was then resuspended in 700 µL QIAzol for RNA analysis.

### Plaque assay

Vero cells (WHO, Geneva, Switzerland) were seeded at a density of 5 x 10^5^ cells per well in 6-well plates. The next day, the medium was removed from the monolayers and 200 µL of 10-fold serial dilutions of virus-containing supernatant in DMEM containing 2% FBS added to respective wells in duplicate. Vero cell monolayers were incubated at 37°C for 2 h, with rocking at 15 min intervals. Monolayers were overlaid with 1% low-melting point SeaPlaque® agarose (Lonza), set at 4°C for at least 20 min, and then returned to the 37°C incubator. Plaques were visualized and counted 4 days later by crystal violet staining.

### NanoString nCounter Assay and Gene Analysis

The NanoString nCounter platform (NanoString, Seattle, WA, USA) was used to quantify mRNA counts in PBMC processed from whole blood at pre- and post-SIV and ZIKV infection time-points. RNA was isolated from cryopreserved PBMC samples collected at pre-SIV inoculation (Wk -3), 2 and 6 weeks post-SIV using the miRNeasy Mini Kit (QIAGEN). RNA was isolated from 1-2 x 10^6^ PBMC resuspended in 700 µl QIAzol collected at week 7/Day -14 pre-ZIKV inoculation, and at days 2, 4, 7, 10, 14 and 21 post-ZIKV challenge using the miRNeasy Micro Kit (QIAGEN). From each sample, 100 ng RNA was loaded in accordance with manufacturer’s instructions for targeted expression with 2 custom-built curated NanoString Human Panels of 44 and 60 genes of interest which both represent gene biomarkers of innate immune activation and response, interferon response, and inflammatory response. Due to the small number of genes represented on the Code Set, nCounter data normalization was performed using a method which calculates a ratio between genes of interest to housekeeping genes(32). The ratio is calculated by dividing the counts of the genes of interest by the geometric mean of 4 housekeeping genes which have the lowest coefficient of variance across all samples. This is done for each sample independently, which generates normalized expression for the genes of interest. Significant differences (nominal *P*-val <0.01) were determined between baseline and infection time points for each group and between groups at each time point.

### Histology

At necropsy representative samples of all tissues and organs were collected in formalin and after fixation were paraffin embedded and sectioned at 3-5 μm. For basic histology, sections were stained with hematoxylin and eosin. All histological findings are summarized in

## Supplemental Table 2

### Immunophenotyping

Isolated PBMC, rectal and peripheral lymph node biopsy cells were assessed for viability with a live/dead stain (Life Technologies) and stained with a panel of antibodies, details described in **Supplementary Table 3,** in brilliant stain buffer (BD Biosciences) to identify innate immune cells as previously described(24). Paraformaldehyde fixed cells were acquired on a LSRII (BD Biosciences) using FACS Diva software (version 8). Samples were analyzed using FlowJo software version 10.8.1 (FlowJo, LLC). All events were first gated on FSC singlets, CD45^+^ leukocytes, live, and then cells according to FSC-A and SSC-A profiles. Immune cells were identified as follows: plasmacytoid dendritic cells (DCs) (CD20^-^CD3^-^HLA-DR^+^CD14^-^ CD123^+^CD11c^-^), myeloid DCs (CD20^-^CD3^-^HLA-DR^+^CD14^-^ CD123^-^CD11c^+^), monocytes (CD20^-^ CD3^-^HLA-DR^+^CD14^+^CD16^+/-^), and neutrophils (CD3^-^CD11b^+^CD14^+^SSC-A^Hi^). AXL positive cells were identified after FMO subtraction and meeting a cellular threshold (≥100 cells/gate).

### Multiplex Bioassay

Cytokine and chemokine levels in plasma and cerebrospinal fluid (CSF) were analyzed using a custom nonhuman primate ProcartaPlex 24-plex immunoassay (ThermoFisher Scientific), per the manufacturer’s protocol. The levels of the analytes were assessed on a Bio-Plex 200 system (Bio-Rad) and analyzed per the manufacturer’s protocol.

### ZIKV RNA Quantification

Viral RNA load was assessed in plasma, CSF, rectal cytobrush supernatant, and tissues using a ZIKV-specific RT-qPCR assay, as previously described(24). RNA was isolated from plasma and rectal cytobrush supernatant collected pre-challenge and at 1, 2, 3, 4, 7, 10, 14 and 21 days post-infection (dpi) and at necropsy (24-28 dpi). RNA was isolated from CSF collected pre-challenge and at days 4, 7 and 21 post-challenge and at necropsy (24-28 dpi). Rectal and PLN biopsy tissues were collected pre-challenge and at days 7 and 21 post-challenge and lymphoid and gut tissues collected at necropsy (24-28 dpi). The iScript Select cDNA Synthesis Kit (Bio-Rad) was used for gene-specific cDNA synthesis and cDNAs were quantified on a QuantStudio Real-Time PCR System (ThermoFisher Scientific). Ct values <39 in at least 2 of the triplicates and falling within the standard curve determined from diluted known quantities of ZIKV genome were considered positive.

### Gut integrity and neuroinflammation

Plasma and/or CSF quantification by ELISA of human soluble CD14 (sCD14), human fatty acid binding protein 2 (FABP2) (Fisher Scientific, Waltham, MA) or human LPS binding protein (LBP) (Biometec, Germany) was performed per the manufacturer’s instruction. Plasma was diluted as follows: 1:200 (sCD14), 1:2 (FABP2), or 1:3 (LBP). CSF was diluted 1:5 (sCD14). Results were analyzed using Prism version 8.4.3 (GraphPad) and using a four- or five-parameter logistic (4- or 5-PL) function for fitting standard curves.

### Anti-ZIKV IgG Quantification

NHP sera and/or plasma samples were assessed for anti-ZIKV envelope (E) IgG binding titers by an Enzyme-Linked Immunosorbent Assay (ELISA). Purified NHP IgG (MyBioSource MBS539659) was serially diluted to establish a range of IgG standards. ZIKV E protein (Fitzgerald Industries International, 30-1932) was diluted to 0.5 μg/mL and was used as the capture antigen. Capture antigen and IgG standards were coated overnight on high-binding 96-well plates (Costar 3590) to produce test and standard wells, respectively. All wells were subsequently blocked in blocking buffer (5% w/v nonfat dried milk (Bio-Rad Laboratories 1706404) and 0.05% v/v Tween-20 in PBS). Samples were diluted 1:100, 1:200, and/or 1:400 and tested in triplicate. NHP IgG standard and test wells were probed by a goat anti-monkey IgG antibody conjugated to Horseradish peroxidase (HRP) (Abcam ab112767). SureBlue Reserve TMB substrate (KPL) was added to all wells to initiate a color change reaction catalyzed by HRP. Reaction was stopped after 30 minutes with 1N HCl (VWR) and absorbance at 450nm (Abs_450_) was measured on an EMax plate reader (Molecular Devices). Abs_450_ of standard wells were used to produce a 5PL logistic fit (GraphPad Prism). Abs_450_ of test wells were converted to μg/mL of anti-ZIKV E IgG binding titers via the 5PL logistic fit.

### Plaque reduction neutralization test (PRNT)

NHP sera collected pre-challenge (Day -14) and at necropsy (24-28 dpi) were tested in PRNT assay for neutralizing antibody production, as previously described(33). The PRNT assay was performed using serial two-fold dilutions of the serum samples. The highest serum dilution reducing plaque numbers by 50% (PRNT_50_) were determined with a limit of detection (LOD) of 1:50. The assay was repeated twice in triplicate using the ZIKV Brazil 2015 virus.

### Statistical analysis

Non-parametric statistical methods were employed for all comparisons, unless otherwise noted. Specifically, Kruskal-Wallis tests were used for comparisons across timepoints in *in vitro* experiments, paired Wilcoxon tests were used to evaluate cell fraction differences to baseline at each timepoint, and Mann-Whitney tests were used to compare continuous values across groups. All analyses were conducted using two-sided tests at the 0.05 level. Analyses were conducted in Prism version 8.4.3h (GraphPad). Significant differences (nominal *P*-val <0.01) in gene expression were determined using a t test that compared baseline and infection time points for each group and between groups at each time point.

### Data Availability

The data that support the findings of this study are available from the corresponding author upon reasonable request. The NanoString nCounter analysis code is available at https://github.com/galelab/OConnor_SIV-ZIKV_coinfection.

## Acknowledgments

The authors thank all members of the Fuller and Gale labs, A. Gustin, R. Ruiz, E. Broderick, J. Brenchley, and A. Huynh for their technical support and helpful discussions and to B. Keele for generously providing the SIVmac239M. We thank the NIAID DAIDS Nonhuman Primate Core Virology Laboratory (NHP CVL) and the Washington National Primate Research Center (WaNPRC), Virology and Immunology Core (V&IC), Seattle Genomics, and University of Washington Research Testing Services (RTS) for assistance with standard assays. We thank the WaNPRC animal staff for the excellent care of the animals. Pictorial illustration was created with biorender.com.

## Funding

This work was funded by National Institutes of Health (NIH)/NIMH K01MH123258 (MAO), University of Washington/Fred Hutchinson Center for AIDS Research iCFAR Award (MAO, Parent Grant: NIH P30AI027757), University of Washington Sexually Transmitted Infections Cooperative Research Center (STI CRC) Developmental Research Project Award (MAO, Parent Grant: NIH/NIAID U19AI113173), and the Washington National Primate Research Center (WaNPRC)/Institute for Translational Health Sciences (ITHS) Ignition Award (MAO, Parent Grants: NIH/ORIP P51OD010425 and National Center for Advancing Translational Sciences (NCATS) (UL1TR000423), (MG, AI145296), and the NIAID DAIDS Nonhuman Primate Core Functional Genomics Laboratory for AIDS Vaccine Research and Development contract (MG, HHSN272201800003C) for infrastructure. Funding for Seattle Genomics is supported in part by the National Institutes of Health, Office of the Director P51OD010425 (Seattle Genomics, MG) and the WaNPRC breeding colony is supported by U42OD011123. The content is solely the responsibility of the authors and does not necessarily represent the official views of the funders. The funders had no role in study design, data collection and analysis, decision to publish, or preparation of the manuscript.

## Author Contributions

JTG, MAO, MG, DHF, and MAO designed and coordinated the studies. TBL, KV, SN, KH, EB, AH, and MAO led the immunological assays and analysis. JTG, KV, SN, PK, and MAO led the virologic assays and analysis. JTG, LSW, and JD led the transcriptomic data generation and analysis. NI, CA, SW, RM, and KAG led specimen collection and RM and KAG led the clinical care of the animals. PTE assisted with the statistical analysis. JTG and MAO led the studies, interpreted the results, and wrote the paper with all co-authors.

## Supporting information captions

*Supporting Information File 1*. Animal characteristics, SIV plasma load, and peripheral CD4 counts

*Supporting Information File 2*. Gross and histopathologic assessment

*Supporting Information File 3*. Primary antibodies used for immunophenotyping analysis

*Supporting Information File 4*. Post-SIV PBMC gene expression for genes shown in **Figure 1D**

*Supporting Information File 5*. Post-ZIKV PBMC gene expression for genes shown in **Figure 5A**

*Supporting Information File 6*. PBMC gene expression kinetics for SIV+ZIKV+ and SIV-ZIKV+ groups shown in **Figure 5C**

*Supporting Information File 7*. ZIKV RNA quantitation in longitudinal samples

*Supporting Information File 8*. ZIKV RNA quantitation in necropsy samples

## Notes

### Competing Interest Statement

The authors have declared no competing interest.

